# Brain activation by a VR-based motor imagery and observation task: An fMRI study

**DOI:** 10.1101/2022.12.05.519096

**Authors:** João D. Nunes, Athanasios Vourvopoulos, Diego Andrés Blanco-Mora, Carolina Jorge, Jean-Claude Fernandes, Sergi Bermudez i Badia, Patrícia Figueiredo

## Abstract

Training motor imagery (MI) and motor observation (MO) tasks is being intensively exploited to promote brain plasticity in the context of post-stroke rehabilitation strategies. The desired brain plasticity mechanisms may benefit from the use of closed-loop neurofeedback, embedded in brain-computer interfaces (BCIs) to provide an alternative non-muscular channel. These can be further augmented through embodied feedback delivered through virtual reality (VR). Here, we used functional magnetic resonance imaging (fMRI) to map brain activation elicited by a VR-based MI-MO BCI task called NeuRow and compared with a conventional non-VR, and MI-only, task based on the Graz BCI paradigm. We found that, in healthy adults, NeuRow elicits stronger brain activation when compared to the Graz task, as well as to an overt motor execution task, recruiting large portions of the parietal and occipital cortices in addition to the motor and premotor cortices. In particular, NeuRow activates the mirror neuron system (MNS), associated with action observation, as well as visual areas related with visual attention and motion processing. We studied a cohort of healthy adults including younger and older subgroups, and found no significant age-related effects in the measured brain activity. Overall, our findings suggest that the virtual representation of the arms in a bimanual MI-MO task engage the brain beyond conventional MI tasks, even in older adults, which we propose could be explored for effective neurorehabilitation protocols.

## Introduction

Stroke is a leading cause of adult long-term disability [1]. Despite significant progress being made in post-stroke rehabilitation, there is still the need for further improvement of current rehabilitation strategies and their outcomes, as most survivors must cope with some degree of disability or loss of independence in activities of daily living (ADL) [2, 3]. Recovery after stroke implies reorganization of the cortex to compensate for the lesioned area. This is possible through neuroplasticity mechanisms, whereby the brain learns and reorganizes itself to compensate for lost functions [4]. Unfortunately, a considerable fraction of patients suffering from strokes affecting their motor function cannot fully benefit from current rehabilitation strategies due to factors such as a low level of motor control, reduced range of motion, pain, or fatigue [5].

Interestingly, research has shown that patients with severe stroke may benefit from motor imagery (MI) and/or motor observation (MO) training through the use of brain-computer interfaces (BCIs). BCI’s can establish an alternative non-muscular channel between the patient’s brain activity and a computer, providing neurofeedback in a closed-loop. This can be used to strengthen key motor pathways that are thought to help promote brain plasticity mechanisms even in the absence of explicit movement [6–9]. The efficacy of such MI/MO BCI training systems for neurorehabilitation strongly depends on the ability of the MI/MO tasks to elicit the desired patterns of motor-related brain activation [10, 11]. In particular, the activation of the mirror neuron system (MNS) would be key to unravel the potential of MI/MO training systems for neurorehabilitation [12–15].

A growing body of research evidences that concurrent MI and MO might be superior to either condition alone in eliciting the desired brain activity [16–20]. However, the optimal type of task and respective instructions for MI / MO interventions remain to be clarified. In particular, some experimental paradigms present conflictsb etween the observed and imagined actions, such that the reported brain activity may include activation related with compensatory mechanisms [20]. Importantly, technological solutions based on virtual reality (VR) are increasingly adopted in post-stroke rehabilitation, and they have the potential to enhance the effectiveness of BCI approaches by providing more ecologically valid feedback on MI/MO performance [21–23].

The neural correlates of MI and MO have been previously investigated, in particular regarding their overlap with the brain regions recruited by motor execution tasks [24]. However, there is some degree of heterogeneity in what regards combined MI-MO tasks, specially considering the wide variety of existing interventional protocols [9]. Importantly, only a few applications rely on ecologically valid scenarios, such as VR. Also, the effects of aging have not been systematically evaluated, which is of particular concern for post-stroke neurorehabilitation applications. This lack of consensus translates into limitations in the design of reliable systems that can effectively be used in rehabilitation.

We have previously developed a VR-based MI-MO task targeting the upper limbs for stroke rehabilitation - NeuRow [25]. It consists in performing MI of rowing using the left or right arm (i.e., MI), while being provided with feedback of task performance through the first-person perspective of the virtual arms of an avatar (i.e., MO). We have shown that such a VR-based task involving the consistent combination of MI and MO may be more powerful than conventional abstract MI tasks such as the ones based on the Graz-BCI paradigm [6, 23, 26]. However, the underlying brain activation remains to be investigated.

Here, we recruit a cohort of healthy adults and use functional magnetic resonance imaging (fMRI) to map brain activation elicited by the NeuRow MI-MO task and compare it with a commonly used abstract MI task based on the Graz BCI paradigm. We aim to 1) map task-specific brain activation patterns; 2) evaluate differences in brain activation patterns between tasks; and 3) determine age-related differences in such brain activation patterns.

## Materials and methods

### Participants

Two groups of healthy right-handed participants were recruited: 11 young adults (mean age 27 ± 4 years) and 10 older adults (mean age 51 ± 6 years). The elderly population was recruited to age-match the stroke population in terms of age-range demographic [27]. The experimental protocol was designed in collaboration with the local healthcare system of Madeira, Portugal (SESARAM), in accordance with the 1964 Declaration of Helsinki, and approved by the scientific and ethic committees of the Central Hospital of Funchal with approval reference number: 21/2019. A written informed consent was obtained from each participant upon recruitment.

### Image Acquisition

Imaging was carried out on a 3T GE Signa HDxt MRI scanner (General Electrics Healthcare, Little Chalfont, United Kingdom) using a 12-channel head coil. fMRI data were acquired using a multi-slice 2D gradient-echo EPI sequence (TR/TE = 2500/30 ms, voxel size = 3.75×3.75×3.00 mm^3^, flip angle = 90°, and FoV = 240×240 mm^2^). For co-registration purposes, whole-brain structural images were also acquired using a T1-weighted 3D Fast Spoiled Gradient-Echo (FSPGR) sequence (TR/TE = 7.8/3.0 ms, voxel size = 1.00×1.00×0.60 mm^3^).

### Experimental Paradigm

The following three tasks were performed for left and right arm movement separately, yielding a total of six fMRI runs (pseudo-randomized order): (a) NeuRow task: motor imagery with motor observation through VR scenario (MI-MO); (b) Graz task: motor imagery only with abstract instructions (MI), (c) and motor execution (ME): finger-tapping. Each fMRI run consisted of 8 trials, each with 20 s of baseline followed by 20 s of task (total run duration 5.33 min) (Fig 1).

**Fig 1.**
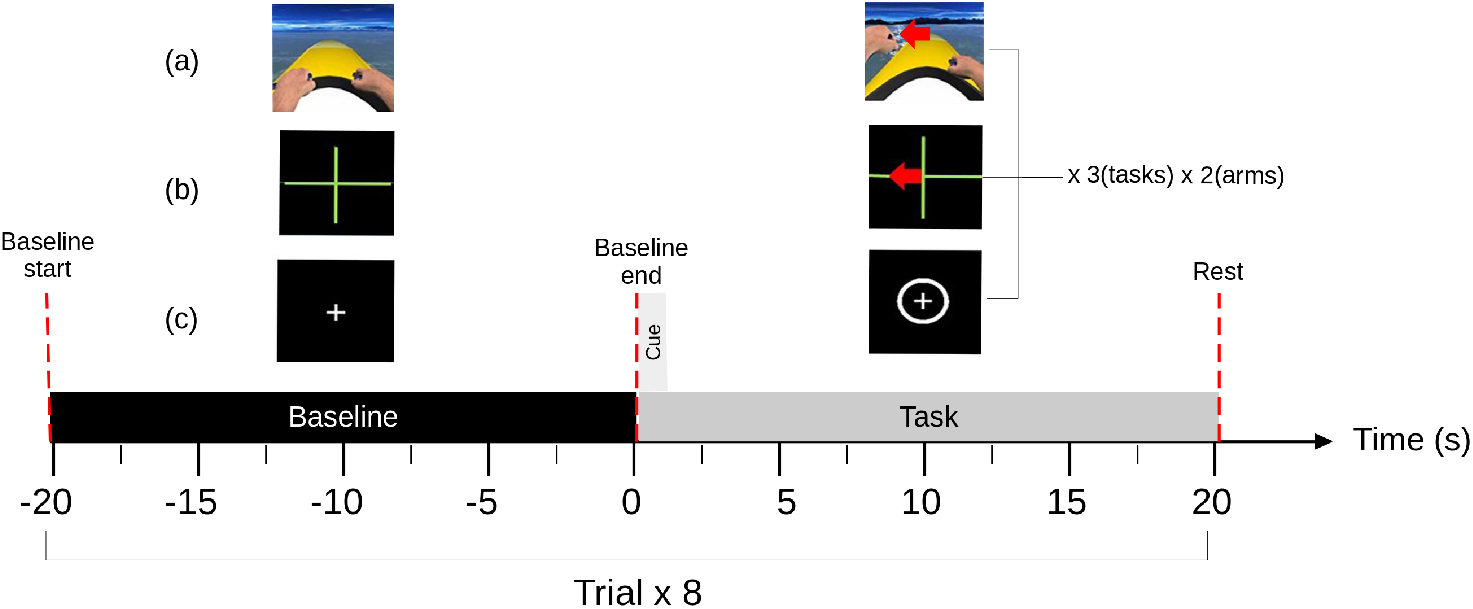
Experimental paradigm. The structure and timings of the trials are shown in the bottom, and the visual instructions are illustrated in the top for each of the three tasks executed by the participants: (a) motor imagery and observation with VR (NeuRow): VR scenario showing own arm movement to cue for left or right arm motor imagery while also providing motor observation. (b) abstract motor imagery (Graz): imagery of arm movement when cued by directional arrow indicating left or right. (c) motor execution (ME): finger tapping when the concentric circle appears around the fixation cross.

For the NeuRow task (MI-MO), participants were instructed to imagine the kinaesthetic experience of rowing. An adaptation of the originally proposed VR training paradigm [25] was used, whereby participants observed the virtual avatar arm moving while instructed to imagine the movement, but did not of control it with their brain activity in a closed-loop. The Graz task (MI) was based on the conventionally used Graz-BCI paradigm [6]. Similar to the NeuRow task, it consisted in imagining the kinaesthetic experience of rowing. However, in this case, a simple directional arrow served as cue (left or right) against an empty black background on the screen (see Figure 1(b)). The Motor Execution (ME) task consisted in a finger tapping task cued using the appearance of a circle outside a fixation cross 1(c)).

### Image Analysis

The fMRI data were analysed using FSL tools (https://fsl.fmrib.ox.ac.uk/fsl). Standard pre-processing was performed including: non-brain tissue removal using FSL’s BET; motion correction FSL’s MCFLIRT, spatial smoothing using a Gaussian kernel with full width at half maximum (FWHM) of 6.5625 mm, and high-pass temporal filtering with a cut-off period of 100 s. Finally, functional images were normalized to the standard Montreal Neurological Institute (MNI152) T1-weighted image (2×2×2 mm^3^ voxel size) by linear registration using FSL’s FLIRT.

Pre-processed fMRI data were then submitted to a first-level general linear model (GLM) analysis. To obtain the explanatory variable (EV) of interest, a boxcar function describing the task paradigm was convolved with the Canonical (Double-Gamma) Haemodynamic Response Function (HRF). Additionally, the 6 motion alignment parameters (3 rotations and 3 translations of the head along the three main axis) were included as confound EV’s of no interest. This GLM was fitted to the data using FSL’s Improved Linear Model (FILM) with pre-whitening to correct for temporal autocorrelations. Contrast of parameter estimate (COPE) maps were obtained for the EV of interest in each subject.

### Statistical Analysis

Group analysis was then performed using a higher-order mixed effects GLM. We followed a backward stepwise regression approach, by first including the group as a factor (young vs. old). Since no effects of group (age) were found, this factor was then removed and the two groups were merged together for further analysis. To identify the group average brain activation pattern associated with each task, one-sample t-tests were performed for each task and arm (ME Right and Left, Graz Right and Left, and NeuRow Right and Left), for the whole subject cohort. A 2-way repeated measures ANOVA was used to assess the effects of task (ME vs. Graz vs. NeuRow) and arm (Left vs. Right), as well as their interaction, again for the whole subject cohort. Since a significant effect of task was found, with no interaction with arm, two-sample paired t-tests were then performed for each pair of tasks (ME vs. Graz, ME vs. NeuRow, and Graz vs. NeuRow), combining right and left arms. In each case, the resulting z-statistic parametric maps were subjected to family-wise thresholding (voxel z>3.1, cluster p<0.05) to yield maps of significant brain activation differences. Finally, we used the probabilistic Harvard-Oxford Cortical [28–31] and Juelich Histological [32–34] atlases to identify brain regions kindred to each activation cluster.

## Results

### Average brain activation for each task and arm

The group average brain activation maps obtained for each task and arm (ME Right and Left, Graz Right and Left, and NeuRow Right and Left) are shown in Figure 2. The brain activation clusters identified in each map are described in Table 1, including the identified brain areas, as well as their volume and peak activation z-stat value and MNI coordinates. There is consistent recruitment of motor and premotor areas across the three tasks. For ME, both the primary motor and premotor cortices, as well as the cerebellum, are clearly activated. For Graz and NeuRow, the premotor cortex is most consistently activated, as well as the putamen. Compared with both ME and Graz, NeuRow recruits additional regions, namely the occipital and parietal cortices and the inferior frontal gyrus, yielding an overall greater volume of brain activation. In terms of the arm side, the expected lateralization of brain activity is observed, with greater activation of the contralateral hemisphere. The lateralization is greater for the right relative to the left arm tasks, also as expected. The observed lateralization patterns are less clear for the Graz task when compared with the ME and NeuRow tasks.

**Fig 2.**
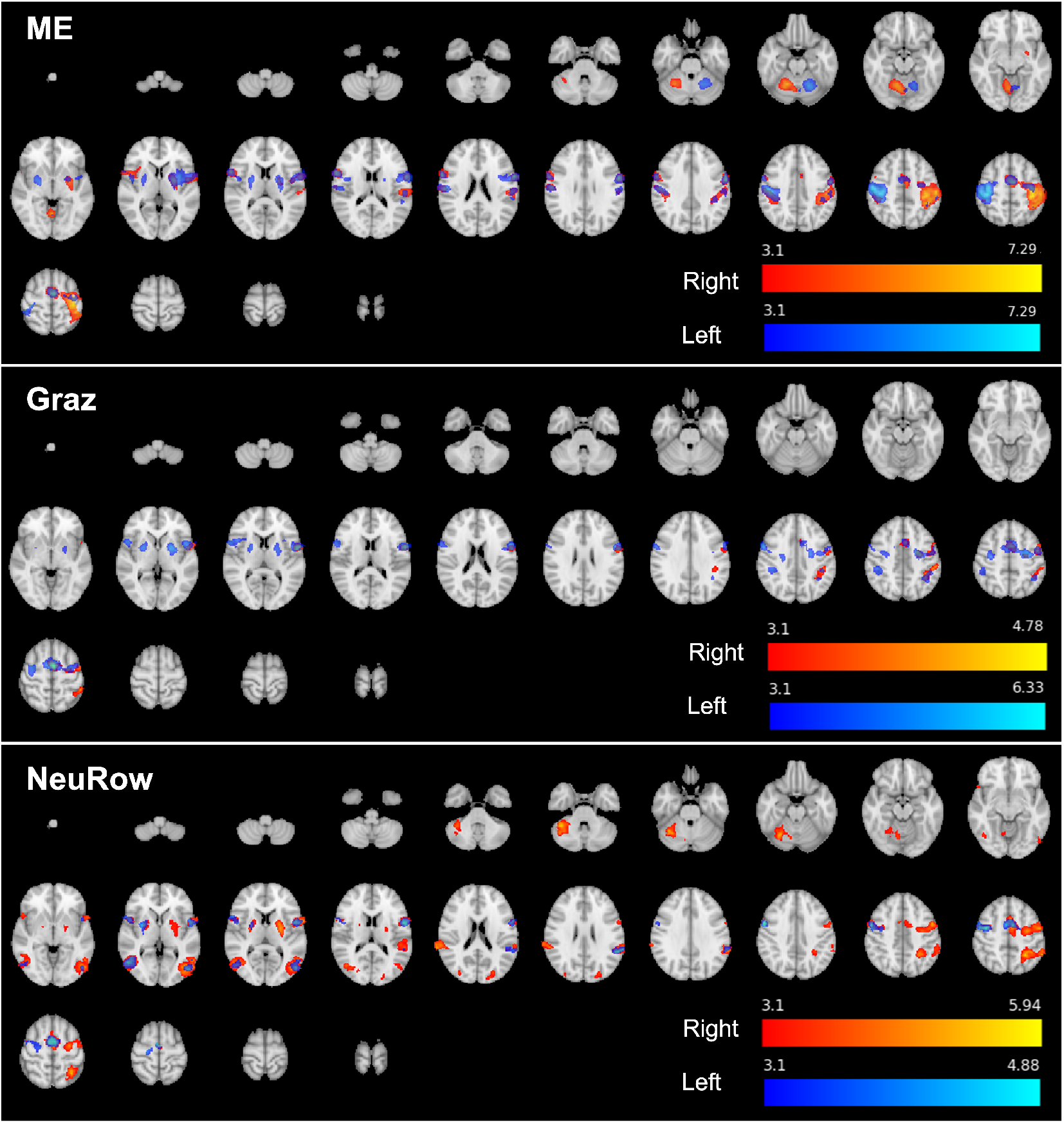
Maps of group activation for each task (ME, Graz and NeuRow) and for each arm (Right and Left), across both age groups (Young and Old). Thresholded z-stat maps (colour) are overlaid on the MNI152 T1-weighted image for a series of representative transverse slices. The brain regions identified in each maps are described in Table 1.

**Table 1.** Clusters of group activation for each task (ME, Graz and NeuRow) and for each arm (Right and Left), across the whole subject cohort (Young and Old).

| Brain Region |  | Volume (voxels) | Max Z-Score | MNI Peak Coordinates (mm) |  |  |
| --- | --- | --- | --- | --- | --- | --- |
|  |  |  |  | X | Y | Z |
| ME Right |  |  |  |  |  |  |
| 1 | Postcentral Gyrus (Right) | 2838 | 7.22 | 38 | -24 | 50 |
| 2 | Precentral Gyrus (Left) | 2525 | 5.66 | -58 | 02 | 18 |
| 3 | Cerebellar Lobule VI (Left) | 847 | 5.39 | -18 | -58 | -22 |
| 4 | Precentral Gyrus (Right) / Broca's Area BA44 | 842 | 5.41 | 62 | 10 | 18 |
| 5 | Parietal Operculum Cortex (Left) / Secondary Somatosensory Cortex | 710 | 4.14 | -56 | -24 | 18 |
| 6 | Juxtapositional Lobule Cortex/ Premotor Cortex | 548 | 5.88 | -02 | -6 | 60 |
| 7 | Right Pallidum | 463 | 4.69 | 22 | -08 | -04 |
| 8 | Precentral Gyrus (Left) / Premotor Cortex | 367 | 4.93 | -38 | -14 | 60 |
| ME Left |  |  |  |  |  |  |
| 1 | Postcentral Gyrus/ Primary Motor Cortex (Right) | 5677 | 7.22 | -36 | -30 | 52 |
| 2 | Supramarginal Gyrus, anterior division / Inferior Parietal Lobule (Right) | 2151 | 5.7 | 44 | -26 | 36 |
| 3 | Cerebellar Lobule V (Right) | 1363 | 6.01 | 14 | -58 | -18 |
| 4 |  | 663 | 4.98 | -24 | -2 | 4 |
| Graz Right |  |  |  |  |  |  |
| 1 | Juxtapositional Lobule Cortex/ Premotor Cortex | 646 | 4.73 | -2.0 | -4.0 | 60 |
| 2 | Supramarginal Gyrus, Anterior Division/ Inferior Parietal Lobule | 692 | 4.39 | -50 | -38 | 54 |
| 3 | Central Opercular Cortex, Secondary Somatosensory Cortex | 1264 | 4.36 | -56 | 0 | 8.0 |
| Graz Left |  |  |  |  |  |  |
| 1 | Right Putamen | 226 | 4.94 | 24 | 4.0 | 4.0 |
| 2 | Supramarginal Gyrus, Anterior Division (Right) | 332 | 4.22 | 44 | -34 | 42 |
| 3 | Supramarginal Gyrus, Posterior Division | 344 | 4.47 | -40 | -48 | 48 |
| 4 | Left Putamen | 395 | 5.01 | -24 | 2.0 | 2.0 |
| 5 | Precentral Gyrus (Right)/ Premotor Cortex | 1213 | 5.10 | -58 | 4.0 | 36 |
| 6 | Juxtapositional Lobule Cortex/ Premotor Cortex | 3307 | 6.27 | -2.0 | -4.0 | 58 |
| NeuRow Right |  |  |  |  |  |  |
| 1 | Lateral Occipital Cortex, Superior Division (Right) | 191 | 3.99 | -24 | -82 | 22 |
| 2 | Right Putamen | 255 | 4.66 | 26 | -4.0 | 6.0 |
| 3 | Central Opercular Cortex | 280 | 4.28 | 50 | 8.0 | 4.0 |
| 4 | Supramarginal Gyrus, Posterior Division (Right)/ Inferior Parietal Lobule | 364 | 4.99 | 72 | -38 | 20 |
| 5 | Lateral Occipital Cortex, superior division (Left) | 485 | 4.96 | -24 | -8.0 | 8.0 |
| 6 | Precentral Gyrus (Right)/ Premotor Cortex | 525 | 4.74 | 38 | -4.0 | 50 |
| 7 | Precentral Gyrus (Right)/ Broca's Area BA44 | 744 | 5.01 | -58 | 6.0 | 10 |
| 8 | Lateral Occipital Cortex, Inferior Division (Right)/ Visual Cortex V5 | 1150 | 5.08 | 56 | -62 | 4.0 |
| 9 | Right Crus I | 1264 | 5.51 | 36 | -56 | -30 |
| 10 | Lateral Occipital Cortex, Inferior Division (Left) | 1399 | 5.51 | -44 | -74 | 4.0 |
| 11 | Superior Parietal Lobule (Left)/ Precuneus Cortex | 1870 | 5.49 | -28 | -52 | 50 |
| 12 | Juxtapositional Lobule Cortex (Premotor Cortex) | 2169 | 5.88 | -2.0 | -4.0 | 60 |
| NeuRow Left |  |  |  |  |  |  |
| 1 | Right Putamen | 212 | 4.37 | 24 | 6.0 | 2.0 |
| 2 | Inferior Frontal Gyrus, Pars Opercularis (Right) | 280 | 4.04 | 56 | 14 | 2.0 |
| 3 | Lateral Occipital Cortex, Inferior Division (Left) | 311 | 4.16 | -50 | -76 | 6.0 |
| 4 | Supramarginal Gyrus, Posterior Division (Left) | 355 | 4.75 | -56 | -46 | 24 |
| 5 | Lateral Occipital Cortex, Inferior Division (Right) | 455 | 4.32 | 54 | -64 | 2.0 |
| 6 | Precentral Gyrus (Right), Broca's Area BA44 | 521 | 4.84 | -56 | 4.0 | 16 |
| 7 | Juxtapositional Lobule Cortex/ Premotor Cortex | 648 | 5.07 | 6.0 | 0.0 | 54 |
| 8 | Precentral Gyrus (Right)/ Premotor Cortex | 849 | 5.65 | 56 | 2.0 | 38 |

### Differences in brain activation between tasks and arms

The 2-way repeated measures ANOVA yielded significant main effects of task and arm, with no significant interactions between them. The maps of significant differences between pairs of tasks (NewRow vs. Graz, NewRow vs. ME, and Graz vs. ME) are presented in Figure 3. The brain regions identified in each map are described in Table 2, including the identified brain areas, as well as their volume and peak activation z-stat value and MNI coordinates. As expected, the ME task more strongly activated the primary motor and sensorimotor cortices, as well as the cerebellum, when compared to both imagery tasks, Graz and NeuRow. Interestingly, a few areas were also more strongly activated by the Graz and NeuRow tasks than ME. While Graz further activated small areas of the frontal and occipital cortices, NeuRow produced greater activation over large areas across the occipital and parietal cortices. This was also evident when directly comparing NeuRow with Graz.

**Fig 3.**
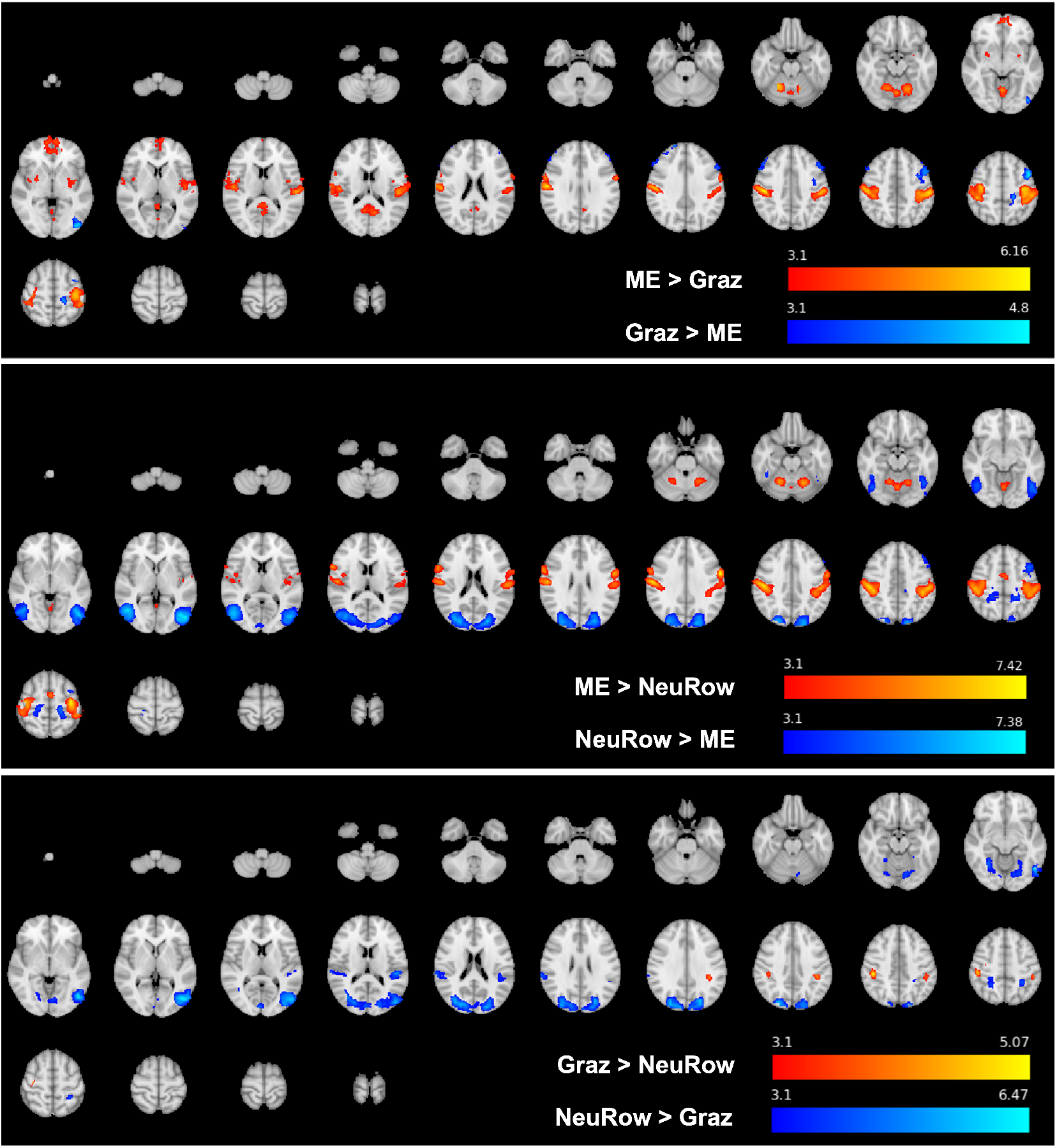
Maps of group activation differences between tasks. Pairwise t-tests between tasks, across both arms (Right and Left) and age group (Young and Old): Graz vs. ME; NeuRow vs. ME; and Graz vs. NeuRow. Thresholded z-stat maps (colour) are overlaid on the MNI152 T1-weighted image for a series of representative transverse slices. The brain regions identified in each maps are described in Table 2.

**Table 2.**
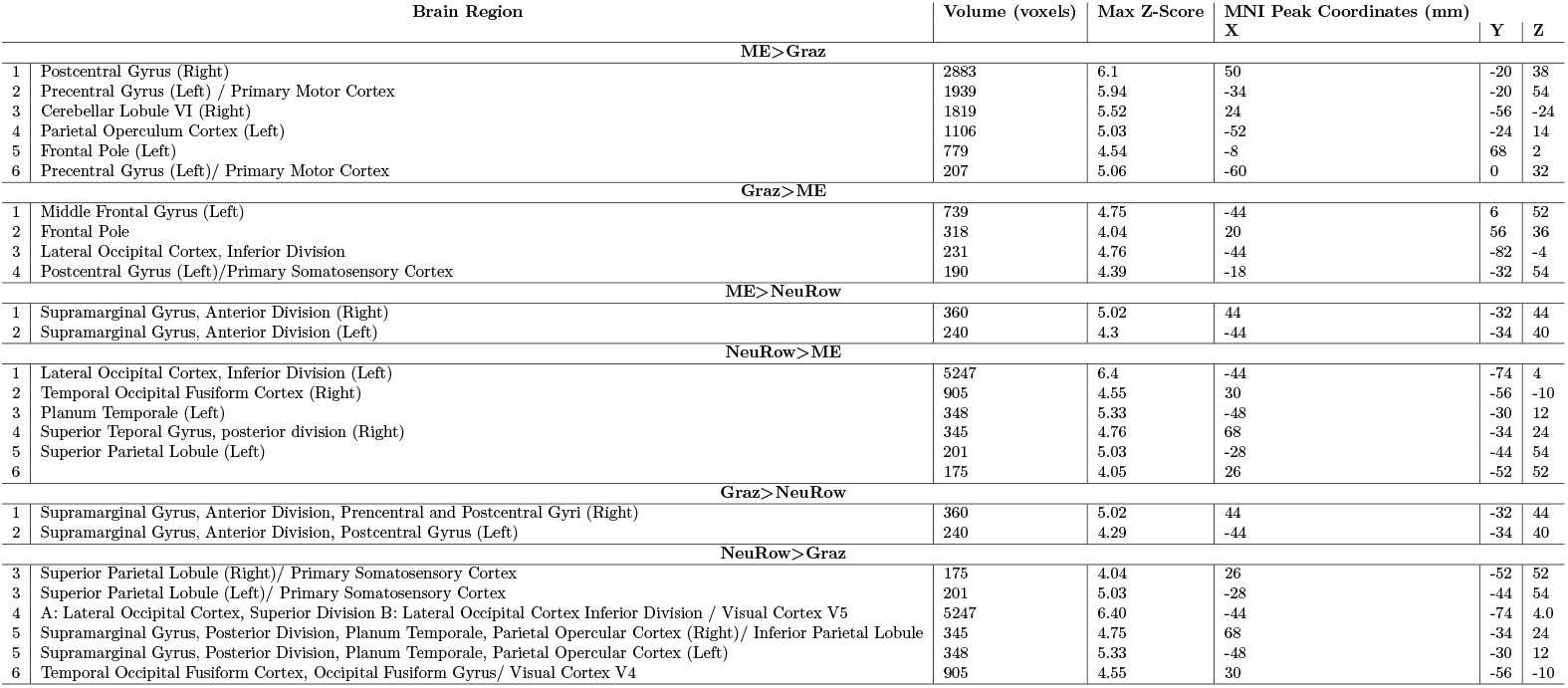
Clusters of group activation differences between tasks. Two-sample paired t-tests between tasks, across both arms (Right and Left) and age group (Young and Old): Graz vs. ME; NeuRow vs. ME; and Graz vs. NeuRow.

## Discussion

In this study, we showed that we could elicit stronger brain activation with our newly developed NeuRow task, combining MI and MO in a VR-like scenario, when compared with a conventional MI task based on the Graz BCI paradigm, which uses abstract instructions and no VR immersion. When compared to the Graz task, as well as to an overt motor execution task, NeuRow recruited a large volume of the brain across the occipital and parietal cortices, additionally to the motor and premotor cortices.

### Recruitment of sensorimotor cortex

The ME task strongly activated the primary motor and somatosensory cortices, as well as cerebellar structures, namely cerebellar lobules I-IV, VI and VIII. There is evidence supporting a topographic organization of the cerebellum, with lobules V-VI and VIII involved in motor processing [35]. Moreover, resting-state functional connectivity studies report correlation of activity in sensorimotor cortices with activity in cerebellar lobules V, VI and VIII [36–38]. Therefore, our findings support that the recruited cerebellar regions are kindred to motor processing, more precisely with a finger tapping movement.

In turn, although with less extension, both imagery tasks, Graz and NeuRow, elicited activity in sensorimotor areas as well, as expected [11]. A more detailed observation shows that the volume recruited by NeuRow and Graz centres mainly in premotor regions, which are typically associated to action preparation [39]. These findings are in line with converging evidence of consistent recruitment of brain regions typically linked to sensorimotor behaviour across ME, MI and MO tasks [10, 24].

In fact, despite the large overall brain activation differences between the NeuRow and Graz tasks, we found no significant differences in the activation of premotor regions. This supports the idea that both tasks are able to effectively promote MI and activate the brain areas involved in action preparation [40]. The fact that these neuronal correlates are also partly shared with the execution of the movement indicates that both MI paradigms are in principle adequate for neurorehabilitation. This is particularly true considering that neuroplastic changes that lead to the recovery of function after stroke are thought to involve the premotor cortex [41].

### Recruitment of additional brain areas

When compared with the Graz task, the NeuRow task further activated areas of the parietal cortex consistent with the MNS [12]. This may be explained by the fact that, unlike Graz, NeuRow involved the observation of the arm movement in addition to imagery. In particular, we found significantly greater activation of the inferior parietal lobule (IPL), namely in its sub-regions PFcm, PF, and PFt [42]. A previous study suggested that the combined activity of these IPL sub-regions with sensorimotor activation may play a specific role in visuospatial and attention-based motor processing [43]. On the other hand, the IPL PFt sub-region has been proposed to be the human homologue of the PF region in primates, which is supposed to contain mirror neurons [24]. These findings corroborate the activation elicited by NeuRow given its motor observation component.

Regarding the greater activation found in the occipital cortex with NeuRow compared with Graz, it is probably the result of a combination of several factors. In general, it is not surprising that greater visual activation is induced by NeuRow given its visual feedback component, which is not present in the Graz task. More specifically, the fact that the visual stimulus consists of a first-person perspective of one’s own arm rowing a moving boat on a lake may explain the activation of area V5 of the primary visual cortex, which is known to be involved in the processing of visual motion [44]. Moreover, we also found greater activation of area V4 of the primary visual cortex, which is thought to integrate information from areas V1 and V2 [45]. However, several studies have shown that V4 neurons may also be directly involved in the processing of a wide range of properties of visual stimuli, including surface properties (colour, shape, texture), movement of the visualized object, and even visual attention [?, 45, 46]. This function is also consistent with the execution of the NeuRow task.

Greater activation of the precuneus was also found. This brain area has been reported to be involved in a wide spectrum of visual tasks, including visuospatial imagery, episodic memory retrieval, and first-person perspective, all of which are present in the NeuRow task [48].

Overall, our findings agree with previous work showing that a combination of MI with MO leads to stronger activation of the brain than either condition alone [17, 18]. Moreover, the realistic virtual environment of NeuRow may contribute to a stronger engagement of various brain areas involved in different aspects of the task, ranging from visual attention to motor preparation and observation. An overall greater engagement of the brain may be desirable in rehabilitation settings, since it is likely associated with improved focus and motivation. In fact, these are essential to enhance adherence to therapy and, thus, rehabilitation outcomes [49]. To confirm our hypothesis, future research should address this question by comparing MI/MO-based BCI-VR strategies with Graz paradigms in post-stroke rehabilitation interventions.

### Age-related effects

Taking in consideration that the NeuRow tool is designed for post-stroke rehabilitation and given that the prevalence of stroke increases with age, it was of the utmost importance to assess whether the observed brain activation patterns changed in an older population. Therefore, we compared the brain activation found in young adults with that of a group of healthy older subjects, with an age range matching the typical age range of stroke patients undergoing neurorehabilitation.

Our analysis did not show any significant differences between the young adults (mean age = 27 ± 4 years) and the older subjects (mean age = 51 ± 6 years). These findings are in accordance with a previous work, showing that MI ability is not affected up to 70 years [50]. Yet, that study reported compromised MI ability in individuals of more than 70 years old. Thus, a different analysis, comparing brain activation in adults less than 70 years old versus older subjects is recommended, as it could provide interesting insights about the question at hand. Indeed, this would be of great importance, since the efficacy of NeuRow and other MI/MO-based BCI paradigms is highly dependable on the individuals’ MI ability.

### Future work

To assess the clinical benefits of the proposed VR-based MI-MO systems over conventional tools, future research should investigate brain activity elicited by these different paradigms in a population of post-stroke survivors undergoing an appropriate rehabilitation intervention. Furthermore, future studies should investigate which of the reported active regions most contribute to the neuroplastic changes that occur after training with the tool. This could allow the design of tailored MI/MO-driven VR BCI systems to achieve greater rehabilitation outcomes.

## Conclusion

We showed that a combined MI-MO task with VR immersion can recruit sensorimotor systems as well as brain structures involved in visual processing and attention-based motor tasks. This work extends on previous research about brain activation during concurrent MI and MO tasks [18, 20, 24, 51], by further including VR and in this way providing more ecological validity. Therefore, although the clinical effects of brain activation after training with NeuRow should be further investigated in a cohort of stroke survivors, we observed in healthy adults an overall stronger brain activation than a conventional abstract MI task and propose that integrating MI and MO in BCI-driven VR into a multidisciplinary stroke rehabilitation program could lead to enhanced recovery.

## Acknowledgments

This work is supported by MACBIOIDI2 (MAC2/1.1b/352), NOVA-LINCS (PEest/UID/CEC/04516/2019), and the Fundação para a Ciência e Tecnologia (FCT) through CEECIND/01073/2018, the LARSyS Project UIDB/50009/2020, the MIGN2Treat Project PTDC/EMD-EMD/29675/2017, and the NeurAugVR Project PTDC/CCI-COM/31485/2017. In addition, the authors would like to acknowledge the contribution of MRI technician, Mr. Sidonio Fernandes, of Hospital “Dr. Nelio Mendoça” in Funchal (Portugal) for supporting the data acquisition.

